# Ketamine Affects Prediction Errors about Statistical Regularities: A Computational Single-Trial Analysis of the Mismatch Negativity

**DOI:** 10.1101/528372

**Authors:** Lilian A. Weber, Andreea O. Diaconescu, Christoph Mathys, André Schmidt, Michael Kometer, Franz Vollenweider, Klaas E. Stephan

**Affiliations:** Translational Neuromodeling Unit (TNU), Institute for Biomedical Engineering, University of Zurich and ETH Zurich, Zurich, Switzerland; Dept. of Psychiatry, University of Basel, Basel, Switzerland; SISSA, Trieste, Italy; Neuropharmacology and Brain Imaging, University Hospital of Psychiatry, University of Zurich, Zurich, Switzerland; Wellcome Centre for Human Neuroimaging, University College London, London, United Kingdom; Max Planck Institute for Metabolism Research, Cologne, Germany

## Abstract

The auditory mismatch negativity (MMN) is significantly reduced in schizophrenia. Notably, a similar MMN reduction can be achieved with NMDA receptor (NMDAR) antagonists. Both phenomena have been interpreted as reflecting an impairment of predictive coding or, more generally, the “Bayesian brain” notion that the brain continuously updates a hierarchical model to infer the causes of its sensory inputs. Specifically, predictive coding views perceptual inference as an NMDAR-dependent process of minimizing hierarchical precision-weighted prediction errors (PEs). Disturbances of this putative process play a key role in hierarchical Bayesian theories of schizophrenia. Here, we provide empirical evidence for this clinical theory, demonstrating the existence of multiple, hierarchically related PEs in a “roving MMN” paradigm.

We applied a computational model (Hierarchical Gaussian Filter, HGF), to single-trial EEG data from healthy volunteers that received the NMDAR antagonist S-ketamine in a placebo-controlled, double-blind, within-subject fashion. Using an unrestricted analysis of the entire time-sensor space, our computational trial-by-trial analysis indicated that low-level PEs (about stimulus transitions) are expressed early (102-207ms post-stimulus), while high-level PEs (about transition probability) are reflected by later components (152-199ms, 215-277ms) of single-trial responses. Furthermore, we find that ketamine significantly diminished the expression of high-level PE responses, implying that NMDAR antagonism disrupts inference on abstract statistical regularities.

Our findings suggest that NMDAR dysfunction impairs hierarchical Bayesian inference about the world’s statistical structure. Beyond the relevance of this finding for schizophrenia, our results illustrate the potential of computational single-trial analyses for assessing potential disease mechanisms.

## Introduction

The auditory mismatch negativity (MMN), an electrophysiological response to rule violations in auditory input streams, has long served as an empirical demonstration that the brain learns the statistical structure of its environment and predicts future sensory inputs (1–3). It plays an important role in psychiatric research, as it fulfils several criteria for a biomarker of schizophrenia (4, 5). Most importantly, a reduction in MMN amplitude is one of the most robust electrophysiological abnormalities in patients with schizophrenia (4–7).

Physiologically, MMN has been shown to depend on intact NMDA (N-methyl-D-aspartic acid) receptor signalling. Following an initial study in monkeys (8), human EEG studies (9–11) using the NMDA receptor (NMDAR) antagonist ketamine also found a significant reduction of MMN responses, although the results show non-trivial variations with ketamine dose, paradigm type and trial definition (12–14). From a neuropharmacological perspective, this renders the MMN paradigm an interesting potential readout of NMDAR function (although with potentially concomitant effects on AMPA receptor function (15)).

The robust impairment of MMN in schizophrenia, and the fact that a similar MMN reduction can be achieved with NMDAR antagonists like ketamine, are in line with the long-standing notion that the pathophysiology of schizophrenia involves NMDAR dysfunction, leading to both cognitive and perceptual abnormalities and positive symptoms (16–22). This has been interpreted as an impairment of perceptual inference under a predictive coding view. In this “Bayesian brain” framework, the brain continuously updates a hierarchical model of its environment to infer the causes of its sensory inputs and predict future events (23–26).

The auditory MMN is believed to reflect such a model update during perceptual inference within the auditory processing hierarchy (3, 27, 28). In particular, in predictive coding, each level of a cortical hierarchy provides predictions about the state of the level below and, in turn, receives a prediction error (PE) signal that reflects the discrepancy between the prediction and the actual state of the level below; this PE signal then serves to update the prediction. This updating process rests on hierarchical message passing between cortical regions, until PEs are minimized on all levels of the hierarchy. While predictions are thought to be communicated by descending (backward) connections, drawing predominantly on glutamatergic NMDAR signaling, sensory PEs are signalled by ascending (forward) connections mainly via glutamatergic AMPA receptors (29). Critically, these ascending PE signals are weighted by the relative precision of bottom-up (sensory) input compared to predictions (priors) from higher levels. The MMN, which is a difference waveform, is then commonly interpreted as the difference in precision-weighted PEs between surprising events (‘deviants’) and more predictable events (‘standards’).

The predictive coding perspective, which understands the MMN as a reflection of perceptual inference in the auditory cortical hierarchy, makes two major predictions:

First, multiple and hierarchically related precision-weighted PEs should underlie the MMN (28). These may become apparent when considering volatility effects during learning (30–32). Volatility determines the learning rate, and even when the real volatility is low or absent in a cognitive paradigm, participants still need to infer the adequate level of volatility as they perform the task. Moreover, in perceptual learning paradigms like MMN, trial-by-trial changes in evoked responses (as measured by EEG) carry information about the temporal dynamics of this learning process (33–38). A suitable model for incorporating volatility in trial-by-trial Bayesian belief updates is the Hierarchical Gaussian Filter (HGF) (30, 39), which quantifies the trajectories of hierarchically related PEs.

Second, the expression of precision-weighted PEs should be sensitive to NMDAR manipulations. According to the framework outlined above, a blockade of NMDARs would lead to a reduction of top-down (predictive) signalling, resulting in less constrained low-level inference about the causes of sensory inputs, and potentially aberrant bottom-up (PE) signalling (19, 20, 40). This could render all events equally surprising and thus blur differences between standard and deviant events (which define the MMN). Such aberrant hierarchical Bayesian inference due to disturbances in the relative weighting of prior beliefs and prediction errors on multiple (sensory and cognitive) levels is at the heart of current computational accounts of schizophrenia (16–18, 22, 41, 42).

A previous MMN study (10) that administered S-ketamine to healthy volunteers focused on the MMN “slope” – the increase of MMN amplitude with the number of standard repetitions, or ‘memory trace effect’. The study demonstrated a reduction of MMN slope at frontal channels under ketamine and interpreted this effect as a disturbance of auditory PE processing. While an important contribution to computational interpretations of MMN, a major limitation of this previous study was the lack of a formal trial-wise model of PEs. Here, we re-analysed this dataset, using a computational single-trial EEG analysis guided by the HGF, to directly test the presence of multiple hierarchically PEs and their susceptibility to NMDA receptor antagonism by S-ketamine.

## Methods and Materials

Details on participants, drug administration, and data acquisition have been provided previously (10, 13); the interested reader is referred to these papers for more information. Here, we only briefly summarize these aspects and focus on the model-based EEG analysis.

### PARTICIPANTS

19 healthy subjects (twelve males, mean age: 26 ± 5.09 years) gave informed written consent and participated in the study, which was approved by the Ethics Committee of the University Hospital of Psychiatry, Zurich. The use of psychoactive drugs was approved by the Swiss Federal Health Office, Department of Pharmacology and Narcotics (DPN), Bern, Switzerland. For further examinations prior to inclusion and additional questionnaire assessments, see (10).

### EXPERIMENTAL PROCEDURE AND DATA PREPROCESSING

The two sessions (placebo and S-ketamine) that all subjects underwent in a counterbalanced fashion were separated by at least two weeks. Both subjects and the experimenter interacting with them were blind to the drug order. For details on the procedure and administration of S-ketamine, please see Supplementary Material.

Electroencephalographic (EEG) activity was recorded during an auditory “roving” oddball paradigm, originally developed by (43) and subsequently modified by (44). The EEG was recorded at a sampling rate of 512 Hz using a Biosemi system with 64 scalp electrodes. Pre-processing and data analysis was performed using SPM12 (http://www.fil.ion.ucl.ac.uk/spm/) and included high- and lowpass filtering and rejection of trials contaminated by eye blinks, as well as bad channels. For details on the paradigm and preprocessing, the reader is referred to the Supplementary Material. The average total number of artifact-free trials was 1211 (sd = 201) in the placebo and 1464.6 (sd = 211.2) in the ketamine condition. The number of artifact-free trials was thus significantly lower in the placebo sessions. However, the resulting non-sphericity was accommodated by our second-level statistical tests (paired t-tests to assess group differences), see Methods section. Note that we did not define categorical events like standard and deviant trials, but instead included all tones in our trial-by-trial analysis.

### MODEL-BASED ANALYSIS

In what follows, we briefly outline our perceptual model before describing the analysis steps used to apply this model to single-trial EEG data. For mathematical details of the model, please refer to the Supplementary Material. In terms of notation, we denote scalars by lower case italics (e.g., *x*), vectors by lower case bold letters (e.g., **x**), and matrices by upper case bold letters (e.g., **X**). Trial numbers are indexed by the superscript (*k*), e.g., *x*^(*k*)^.

### Perceptual Model: The Hierarchical Gaussian Filter (HGF)

To describe a participant’s perceptual inference and learning during this roving MMN paradigm, we use a multivariate version of the Hierarchical Gaussian Filter (HGF), a generic Bayesian model introduced by (30) that has been applied in various contexts, such as associative learning (31, 45), social learning (32, 46), spatial attention (47), or visual discrimination (48).

In the present task, participants were exposed to a tone sequence with 7 different tones. Our modeling approach assumes that in this context, an agent infers two hidden states in the world: (i) the current (probabilistic) “laws” underlying the observed tone statistics – in our case, a matrix **X**_2_ of pair-wise transition probabilities between all tones, and (ii) the current level of environmental volatility, i.e., how quickly the inferred laws seem to change. This is represented in our model by the volatility *x*_3_, which is the degree to which the transition probabilities in **x**_2_ change from trial to trial. The rationale for tracking this quantity is that agents should learn faster – i.e., update their beliefs about the statistical laws in the environment according to prediction errors – if they experience the current environment to be changing rather than stable. Figure 1 shows a visualization of the corresponding generative model.

**Figure 1.**
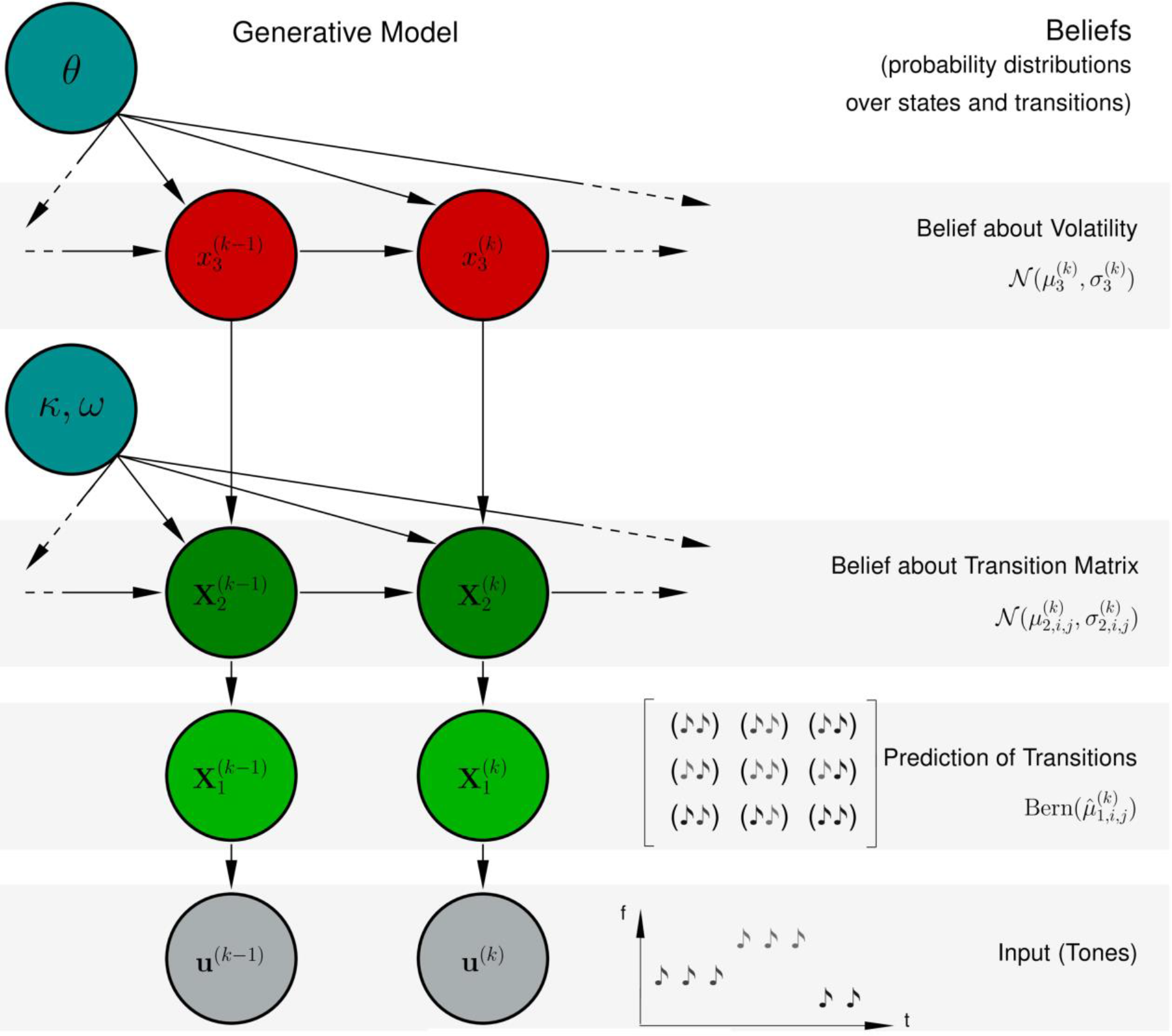
The Perceptual Model: A multivariate version of the binary three-level Hierarchical Gaussian Filter (HGF). The agent infers upon two continuous quantities: The transition tendencies from one tone (frequency) to another, stored in the transition matrix **X**_2_, and the (common) volatility of these tendencies, *X*_3_. To employ this model, the agent only has to follow simple one-step update rules for its beliefs (parameterized by their mean μ and variance σ) about these quantities (updates described in the main text details described in the supplementary material).

On each trial, the agent updates his/her beliefs about these two environmental states, given the new sensory input (i.e., tone). We denote these updated (posterior) beliefs in the following by their mean *μ* and their precision (or certainty) *π* (the inverse of variance, or uncertainty, *σ*). In the HGF, the general form of the update of the posterior mean at hierarchical level *i* on trial *k* is:

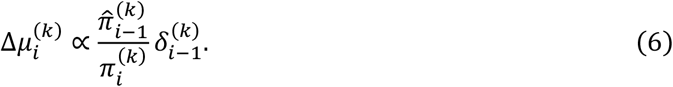

Here, 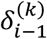 denotes the PE about the state on the level below, which is weighted by a ratio of precisions: 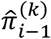 is the precision of the prediction about the level below (*i* − 1), while 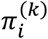 is the precision of the current belief at the current hierarchical level *i*. The intuition behind this is that an agent’s belief updates should be more strongly driven by PEs when the precision of predictions about the input is high relative to the precision of beliefs in the current estimate (e.g., when the environment is currently perceived as being volatile).

The specific update equations for the two levels of our model are given in the Supplementary Material. For a detailed derivation of the update equations and the updates of the precisions, the interested reader is referred to (30). Usually, in the HGF, subject-specific perceptual parameters describe the individual learning style of an agent. Since the current paradigm does not involve behavioral responses to the tones, and thus the model could not be fitted to behavior, we used the parameters (learning rates on both hierarchical levels, and starting values of the beliefs) of a surprise-minimizing Bayesian observer for all participants (for details, see Supplementary Material). The resulting PE trajectories (Figure 2) were subsequently used as regressors in a general linear model (GLM).

### Computational quantities: The precision-weighted prediction errors

The MMN has been interpreted as a precision-weighted PE (or model update signal) during auditory perceptual inference and statistical learning (28, 49, 27, 36, 50, 3, 51). In our model, two hierarchical levels are updated in response to new auditory inputs: the current estimate of the transition probabilities (**μ**_**2**_), and the current estimate of environmental volatility (*μ*_3_). The corresponding precision-weighted PEs driving these updates are hierarchically related and are computed sequentially: the agent first needs to update **μ**_**2**_ (using the low-level PE about **μ**_**1**_) before evaluating its high-level PE with respect to **μ**_**2**_, which is then used to update *μ*_3_.

The questions we address in this paper are whether these precision-weighted PEs, which we denote by *ε*_2_ and *ε*_3_,

i. are reflected by trial-by-trial variations in the amplitude of evoked responses;
ii. their hierarchical relation in the model is mirrored by a corresponding temporal relation in their electrophysiological correlates;
iii. whether NMDAR antagonism by S-ketamine alters the electrophysiological expression of these PEs.

### Single-trial EEG analysis: The General Linear Model

We looked for manifestations of our two computational quantities (*ε*_2_ and *ε*_3_) in the event-related EEG responses for each trial in a time window from 100ms to 400ms post-stimulus. We focused on this time window in order to model learning-induced modulations of both the MMN and the P300 waveforms.

The data from each trial in each session were converted into scalp images for all 64 channels and 91 time points using a voxel size of 4.25 mm × 5.38 mm × 3.33 ms. The images were constructed using linear interpolation for removed bad channels and smoothing to accommodate for between-subject spatial variability in channel space.

Our vectors of precision-weighted PEs served as regressors in a GLM of trial-wise EEG signals for each subject and each session separately, correcting for multiple comparisons over the entire time-sensor matrix, using Gaussian random field theory (52). We did not orthogonalise the regressors. Figure 2 summarizes the analysis steps for the model-based GLM.

**Figure 2.**
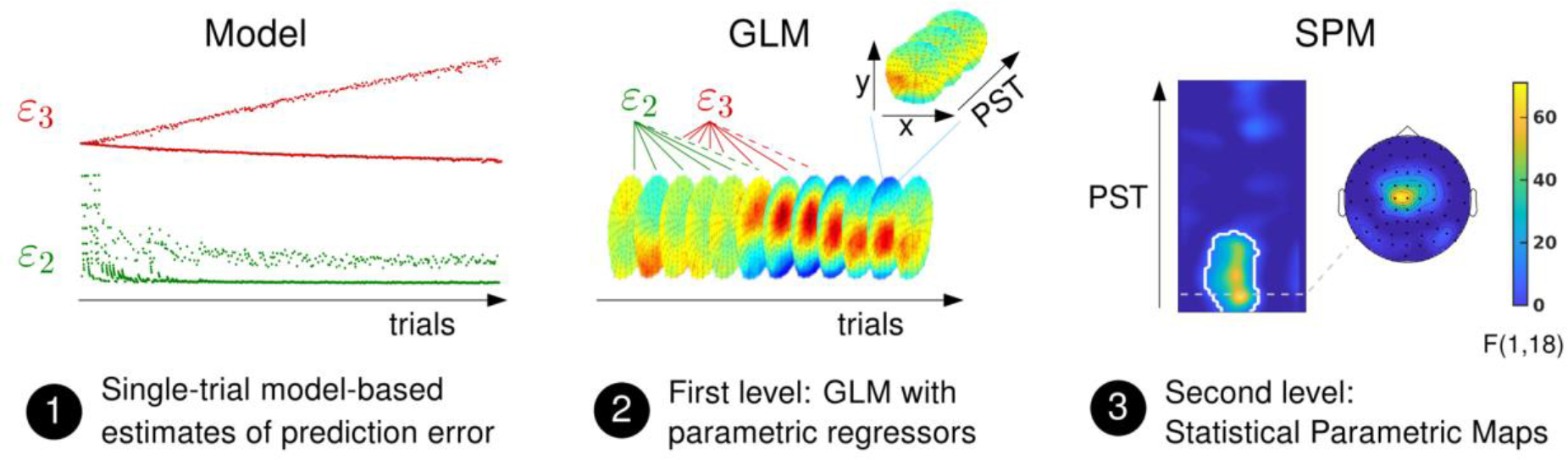
Sketch of the analysis pipeline for the model-based analysis. First, we simulated an agent’s beliefs using our hierarchical Bayesian model, which provided us with an estimate of precision-weighted PE on two hierarchical levels (*ε*_2_ and *ε*_3_) for each trial in each session. At first glance, it may seem that the *ε*_3_ regressor simply amounts to a drift-like signal. This, however, is not the case; the design of our experiment, with prolonged trains of identical stimuli that exchange each other, leads to separate monotonic changes in log-volatility estimates for standard and deviant trials, with jump-like transitions between them (see Figure S1C). Second, we used these estimates as parametric regressors in a GLM of the single-trial EEG signal in each session of each subject separately (peri-stimulus time window of 100 - 400 ms after tone onset) and computed the first level statistics. Third, the beta values for each quantity and each subject in each session entered the second level analysis. We performed random effects group analysis across all 19 participants separately for each drug condition in one-sample T-tests and used F-Tests to examine correlations of EEG amplitudes with our computational quantities of interest, resulting in thresholded F maps across within-trial time and sensor space. PST = peri-stimulus time.

Random effects group analysis across all 19 participants was performed using a standard summary statistics approach (53). We employed one-sample T-tests as second level models, separately for each drug condition, and used F-tests to simultaneously examine positive and negative relations of EEG amplitudes with the trajectories of our computational quantities. To examine differences between the two drug conditions, we tested for reduced responses under ketamine using a paired t-test.

For all analyses, we report any results that survived family-wise error (FWE) correction, based on Gaussian random field theory, across the entire volume (time×sensor space) at the cluster level (p<0.05) with a cluster defining threshold (CDT) of p<0.001 (54). Notably, all reported results also survive whole-volume correction at the peak-level (p<0.05); the associated p-values are not included in the main text but listed in the tables.

## Results

For each computational quantity of interest, our model-based EEG analysis proceeded in two steps: first, we performed whole-volume (spatiotemporal) analyses to search for representations of our quantities in single-trial EEG responses; second, we examined whether these electrophysiological representations of trial-wise PEs differed significantly between ketamine and placebo.

### Low-level precision-weighted prediction errors

By fitting computational trajectories to participants’ single-trial EEG data, we found that under placebo, there was a significant trial-by-trial relation between 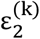 (the precision-weighted transition PE) and EEG activity between 102ms and 207ms post-stimulus, peaking at 121ms at central channels (whole-volume cluster-level FWE corrected, p=2.8e-08, with a cluster-defining threshold (CDT) of p<0.001; Figure 3; Table 1). This time window includes the typical time when the negativity of the roving MMN is observed (43, 44, 49). This suggests that the MMN typically observed in roving MMN paradigms reflects the difference in low-level precision-weighted PEs about stimulus transitions between the subsets of trials labeled as ‘standards’ and ‘deviants’ by the experimenter.

**Figure 3.**
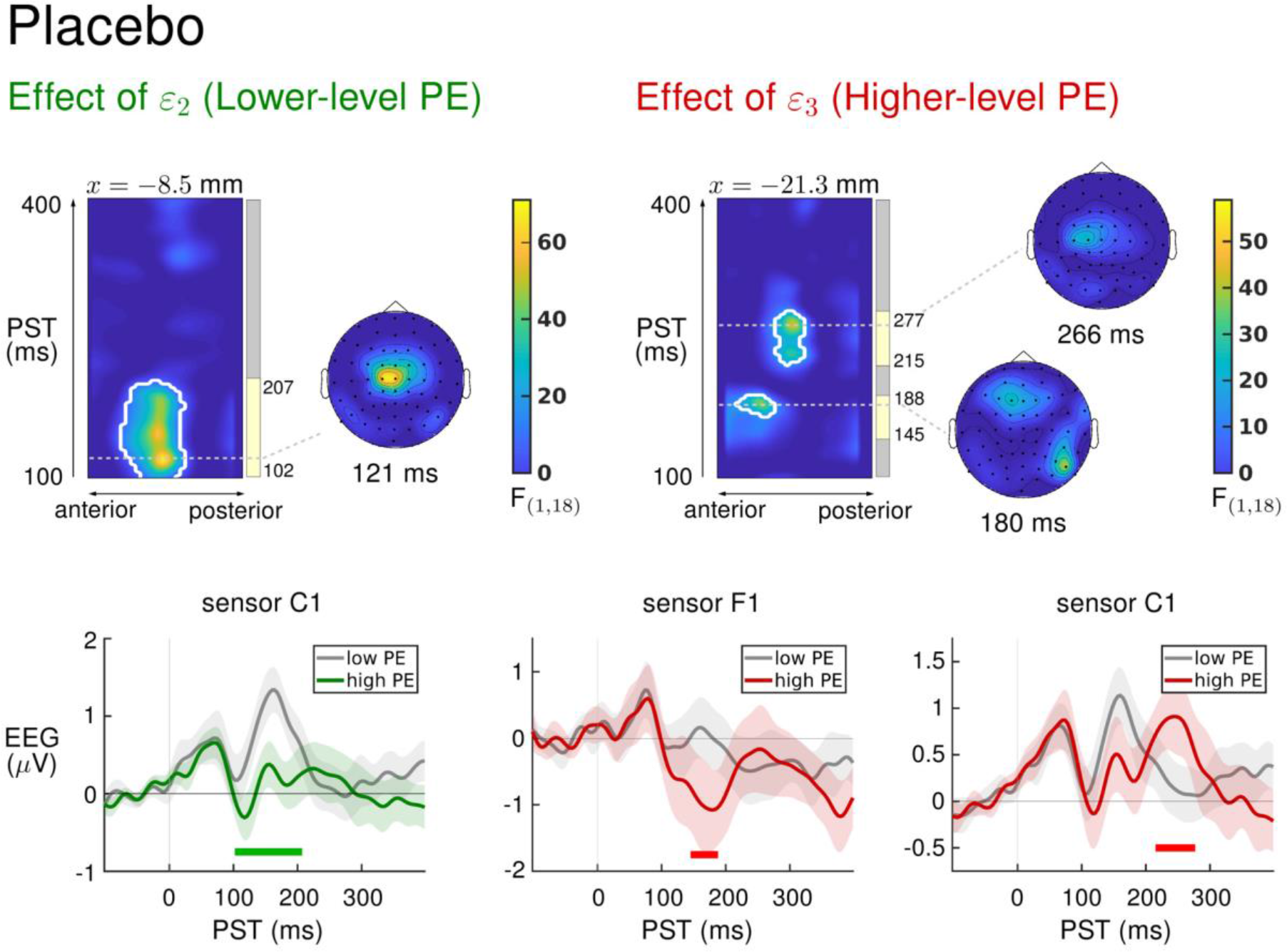
Results of the model-based EEG analysis in the placebo condition: effects of the high- and the low-level PE. *Upper part:* The left side always shows an F-map across the scalp dimension *y* (from posterior to anterior, x-axis), and across peristimulus time (y-axis), at the spatial *x*-location indicated above the map. Significant F values (p < 0.05, whole-volume FWE-corrected at the cluster-level with a cluster-defining threshold of p < 0.001) are marked by white contours. Time-windows of significant correlations are indicated by the yellow bars next to the colored clusters of significant F values. The scalp maps next to the F-maps always show the F-map at the indicated peristimulus time point, corresponding to the peak of that cluster, across a 2D representation of the sensor layout. We found significant correlations of the EEG signal with our two computational quantities across fronto-central and temporal channels. For the lower-level PE, ε_2_, the correlation peaked at 121 ms post-stimulus at central channels, for the higher-level PE, ε_3_, it peaked at 180 ms at frontal channels, at 184 ms at temporal channels (not shown here), and at 266 ms post-stimulus at left central channels. *Lower part:* Average EEG responses to the 10% highest and the 10% lowest PE values at exemplary sensors within significant clusters. Green: High values in low-level PEs correlated with an increased negativity between 102 and 207 ms post-stimulus (sensor C1). Red: High values in high-level PEs correlated with an increased negativity between 145 and 188 ms post-stimulus (sensor F1), and an increased central positivity between 215 and 277 ms post-stimulus (sensor C1).

Under ketamine infusion, we found a similar activation pattern, with significant clusters of activity at fronto-central electrodes between 10ms and 188ms (p=3.1e-08), and at left temporal channels between 105ms and 188ms, peaking at 141ms post-stimulus (p=6.3e-06; Figure S2, Table S3).

**Table 1.**
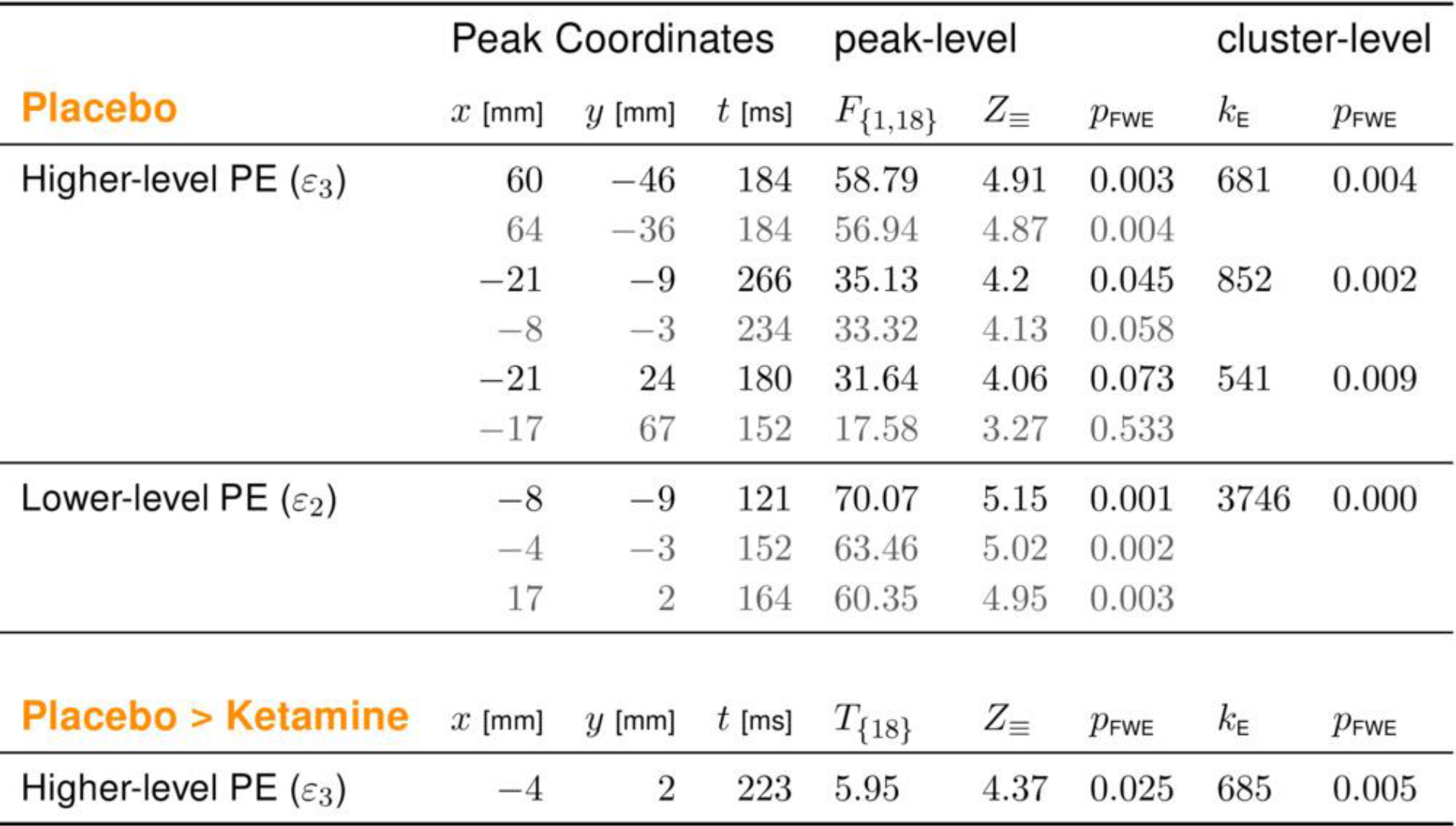
Significant representations of the two computational quantities under placebo, and drug differences for the representation of the higher-level PE ε_3_ in Placebo vs. Ketamine condition. The table lists the peak coordinates, F/t values, corresponding Z values, whole-volume FWE-corrected p-values at the voxel level, cluster size (k_E_) and FWE-corrected p-values at the cluster level of voxels showing significant correlations of EEG signal with the trajectory of one of the precision-weighted PEs ε_3_ and ε_2_ (F-test, p < 0.05 whole-volume FWE-corrected at the cluster-level with a cluster-defining threshold of p < 0.001), and, for the drug difference, of voxels showing significantly stronger representation of ε_3_ under placebo compared to the ketamine condition (paired t-test, p < 0.05 whole-volume FWE-corrected at the cluster-level with a cluster-defining threshold of p < 0.001). No significant drug differences were found for the lower-level PE.

### High-level precision-weighted prediction errors

In the placebo condition, we found a significant trial-by-trial relation between 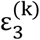 (the precision-weighted PE that serves to update volatility estimates) and EEG activity, both in an early time window (152ms to 199ms, peaking at 184ms at right temporal channels, p=0.004, and from 145ms to 188ms, peaking at 180ms at frontal channels, p=0.009) and in a later time window (between 215ms and 277ms, peaking at 266ms post-stimulus, p=0.002; Figure 3; Table 1), where high-level prediction errors correlated with an increased central positivity corresponding to the P3a component of the auditory evoked potential (55).

Under ketamine, we found a similar relationship of EEG amplitudes with the higher-level PE in the early time window (148ms to 211ms, peaking at 160ms at left temporal channels, p=0.04, and 156ms to 215ms, peaking at 207ms at fronto-central channels, p=0.008), but the later cluster occurred only much later (297ms to 398ms, peaking at 375ms at left temporal channels, p=0.021, and 324ms to 398ms, peaking at 398ms at fronto-central channels, p=0.001; Figure S2, Table S3). While the timing of this late effect is reminiscent of the P3b component, its scalp distribution was not centered on parietal channels, as would be characteristic for P3b (55, 56), but instead looked very similar to the earlier cluster, with a peak at fronto-central channels.

### Effects of ketamine on PE representations

We tested for drug differences in activity elicited by precision-weighted PEs using paired t-tests at the second level. We found no significant differences in activation by 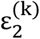 in the ketamine compared to the placebo condition. In contrast, the activation by 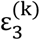 the higher-level PE informing volatility estimates, was significantly reduced under ketamine as compared to placebo in a time window between 207ms and 250ms post stimulus, peaking at 223ms across fronto-central channels (p = 0.005; Figure 4; Table 1). That is, the trial-by-trial relation between EEG signal and the higher-level PE was significantly more pronounced under placebo than under ketamine in this time window.

**Figure 4.**
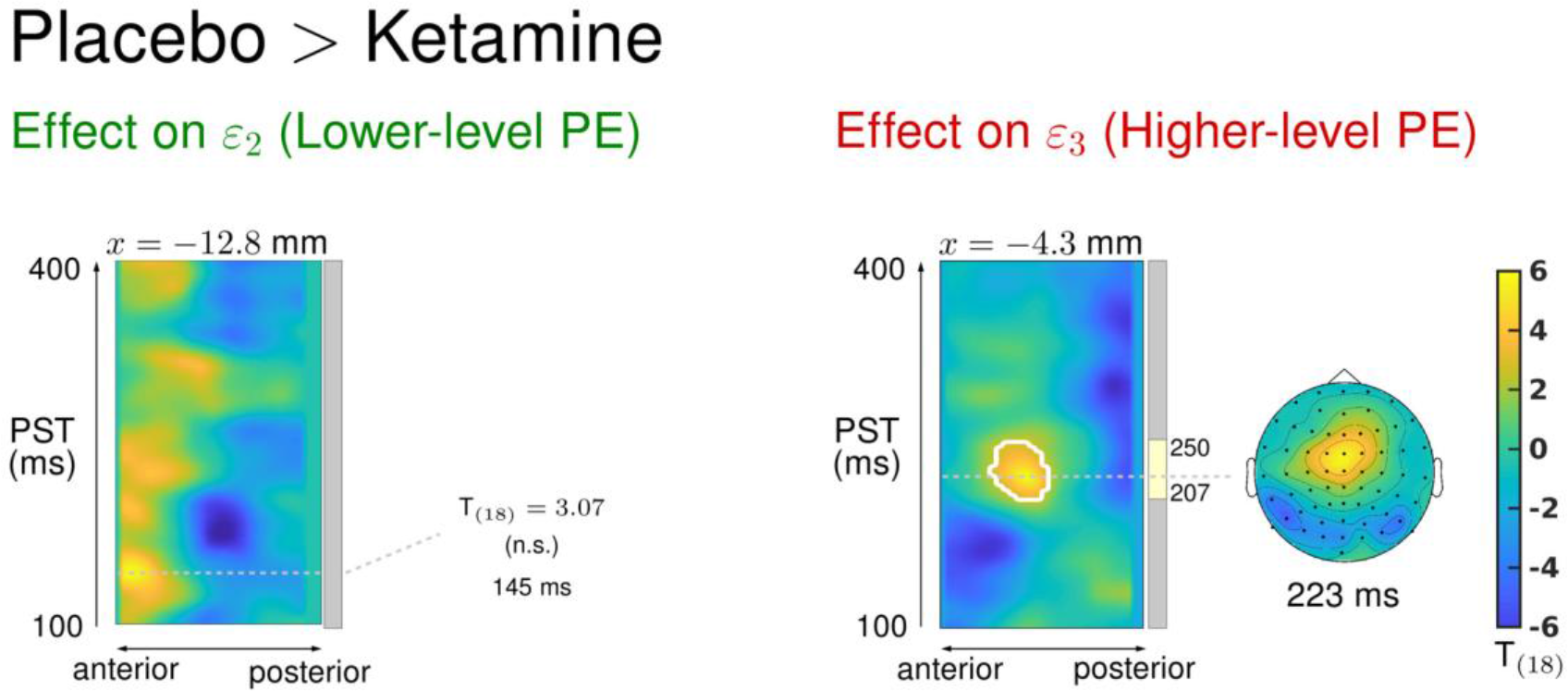
Drug effect on the representation of the lower-level PE (ε_2_, left) and the higher-level PE (ε_3_, right). The left side always shows a T-map for the paired T-test (Placebo – Ketamine) across the scalp dimension *y* (from posterior to anterior, x-axis), and across peristimulus time (y-axis), at the spatial *x*-location indicated above the map. Significant T values (p < 0.05, whole-volume FWE-corrected at the cluster-level with a cluster-defining threshold of p < 0.001) are marked by white contours. Time-windows of significant correlations are indicated by the yellow bars next to the colored clusters of significant F values. The scalp map next to the T-map shows the T-map at the indicated peristimulus time point, corresponding to the peak of that cluster, across a 2D representation of the sensor layout. The effect of the higher-level PE was stronger under placebo compared to ketamine between 207 ms and 250 ms post-stimulus, peaking at 223 ms at fronto-central channels. No significant drug effects were found for the lower-level PE.

## Discussion

Current theories of schizophrenia conceptualize psychotic symptoms as disturbed hierarchical Bayesian inference, characterized by an imbalance in the relative weight (precision) assigned to prior beliefs (or predictions) and new sensory information that elicits PEs (18, 19, 22). Neurobiologically, this disturbance of hierarchical Bayesian inference is thought to result from alterations of NMDAR-dependent synaptic plasticity and to be reflected by abnormalities in perceptual paradigms, such as the auditory mismatch negativity (MMN) (16, 17, 21). Based on a computational single-trial analysis of the MMN under ketamine, the results from the current study are largely supportive of two major predictions: (i) multiple and hierarchically related precision-weighted PEs should underlie the MMN, and (ii) the expression of precision-weighted PEs should be sensitive to NMDAR manipulations.

### MULTIPLE, HIERARCHICALLY RELATED PREDICTION ERRORS UNDERLIE THE MMN

The auditory MMN has been interpreted as reflecting model updates in an auditory processing hierarchy (27, 36, 49). In our Bayesian learning model, levels of a belief hierarchy are updated in response to two different precision-weighted PE signals (30): a low-level PE that quantifies the mismatch between expected and actual tone transitions and a higher-level PE that quantifies the change in estimated uncertainty about transition probabilities in the light of new input and which is used to update estimates of environmental volatility. Effects of volatility on mismatch signals have been reported previously (57–59).

Notably, in the present study, the observed timing of low-level and high-level precision-weighted PE responses under placebo coincided with the timing of MMN and P3a components, respectively, previously shown to reflect related, but dissociable stages of automatic deviance processing (60, 61). Furthermore, the temporal succession of these two PE signatures mirrored the temporal order as predicted by the computational model.

### KETAMINE INTERFERES WITH HIGH-LEVEL BELIEF UPDATES

We found that ketamine changed the electrophysiological expression of the higher-level (but not lower-level) PE. Other authors have reported ketamine-induced changes of the deviant-related negativity at an earlier time corresponding to our lower-level PE representation and the classical MMN latency (9, 10). One difficulty for comparing these reports to the current results is that the timing of ketamine effects in previously reported ERP analyses strongly depended on the type of MMN paradigm, the definition of “standards”, and the choice of electrodes and time windows (9–14). For example, using classical averaging-based ERP analysis restricted to the early MMN time window (100 to 200ms after tone onset) and a subset of fronto-central and temporal channels, Schmidt and colleagues (10) found an attenuation of early MMN amplitudes in frontal channels under ketamine in the same dataset used here. By contrast, our model-based analysis, which considers all sensors and time points under multiple comparison correction, locates the dominant effect of ketamine in the time window of the P3a. This is also consistent with another set of ERP results from the same dataset (13) – where, across all sensors and time points, a significant drug effect was found exclusively in a time window (220-240ms) that was later than the classical MMN latency – and with literature on how ketamine attenuates later ERP components such as the P3 (12, 56, 62).

Our finding that ketamine altered high-level PEs can also be compared to previous dynamic causal modeling (DCM) studies that examined the effects of ketamine during auditory roving MMN paradigms. While these studies (which used different approaches to modeling the input stream) gave different answers, both localized the effect of ketamine at higher levels of the auditory hierarchy. One study found that the effect of ketamine was best explained by changes of inhibition within frontal sources (63). Previous DCM analyses of our own dataset (13) suggested reduced bottom-up connectivity from auditory cortex (A1) to superior temporal gyrus (STG) under ketamine, compatible with disturbed computation of higher-level PEs in STG by impairing message passing from A1.

Interestingly, in our study, the high-level PE showed an effect under ketamine both in early and late time windows (Table 2). While the late effect corresponded to the P3a under placebo, it occurred later under ketamine, around the typical time of ERP components related to conscious processing and context updating (55, 64, 65). Speculatively, this could reflect a less automatic, stimulus-driven processing (as typically associated with P3a) (55, 56, 65) of volatility under ketamine.

It is important to note that our results do not allow for a unique interpretation of ketamine effects in computational terms. If one assumes a strictly monotonic relation between EEG amplitude and PEs, our finding suggests that ketamine reduces learning about environmental volatility. Depending on context, this can both lead to inflated estimates of volatility (slowed representation of stability after periods of inconstancy) or diminished ones (in the opposite case). A previous study using ketamine found reduced stabilization of an internal model of environmental regularities during instrumental learning (66). One may be tempted to interpret this as an overestimation of volatility under ketamine; however, the previous model derived from a different computational concept, making direct comparisons problematic.

## LIMITATIONS

The HGF parameters allow for expression of individual differences in learning (with potential relations to neuromodulatory mechanisms (30, 67)). A main limitation of our approach is that we cannot infer upon such subject-specific learning styles, simply because the MMN paradigm does not provide behavioral responses to which the model could be fitted. Similar to (37), we therefore used the parameters of a surprise-minimizing Bayesian observer for each of the tone sequences and simulated belief trajectories accordingly. An important future extension of HGF applications to MMN paradigms would be the formulation of a forward model from belief updates to EEG signals. This would allow for estimating subject-specific model parameters from single-trial EEG data directly.

A second limitation concerns the relatively small sample size (N=19). This renders it difficult to interpret negative results, such as the lack of ketamine effects on low-level PEs. This will need to be addressed in future studies with larger samples and/or meta-analyses.

## CONCLUSION AND OUTLOOK

This study presents evidence for the role of hierarchically related PEs in the auditory MMN. While ketamine-induced reductions of MMN have been reported previously, our study enables two new insights by taking an explicitly computational perspective and analyzing trial-by-trial belief updates. First, we offer an interpretation of two mismatch-related ERP components, the MMN and the P3a, in terms of hierarchically related PEs that are expressed trial-by-trial and reflect the updating of a hierarchical model of the environment’s statistical structure. Additionally, a reduced expression of the higher-level PE under infusion of S-ketamine suggests a disturbance of high-level inference about environmental volatility by perturbation of NMDA receptors.

Our results are clinically important as they support a bridge between physiology (NMDAR function) and computation (hierarchical Bayesian inference) as proposed by predictive coding theories of schizophrenia. By linking physiological indices of abnormal perceptual inference to their algorithmic interpretation in terms of hierarchically related PEs, the present work provides a starting point for future attempts to understand individual alterations of MMN in schizophrenia mechanistically. We hope that this will contribute to the development of computational assays for improved differential diagnosis and treatment prediction in schizophrenia (68–70).

