## Supplementary Material for "Ketamine Affects Prediction Errors about Statistical Regularities: A Computational Single-Trial Analysis of the Mismatch Negativity"

**Overview:**


- Supplementary Results:
  - Drug administration
  - Paradigm
  - Preprocessing of EEG data
  - Computational model
    - Tab. S1: Priors on perceptual parameters
    - Tab. S2: Comparison of parameter values across drug conditions
    - Fig. S1: Example of simulation results
- Supplementary Results
  - Fig. S2: Results of model-based analysis in ketamine condition
  - Tab. S3: Significant cluster for contrast Placebo > Ketamine on epsilon3, excluding subject 14
  - Tab. S4: Significant cluster for contrast Placebo > Ketamine on epsilon3, excluding four subjects affected by cabling errors

### 1 Supplementary Methods

#### 1.1 Drug administration

S-ketamine was administered using an indwelling catheter that was placed in the antecubital vein of the non-dominant arm. An initial bolus injection of 10 mg over 5 min was followed, after a 1 min break, by a continuous infusion with 0.006 mg/kg per min over 80 min. The initial dose was reduced by 10% every 10 min to keep S-ketamine plasma level fairly constant (1, 2). The procedure in the placebo session was equivalent for administering an infusion of physiological sodium chloride solution and 5% glucose. Each subject was kept under constant supervision until all drug effects had worn off, and was then released into the custody of a partner or immediate relative.

#### 1.2 Paradigm

The E-prime software (3) was used to generate acoustic stimuli that were presented binaurally through headphones.

The stimuli consisted of seamlessly connected trains of pure sinusoidal tones (70 ms duration, 500 ms inter-stimulus interval) with a roving frequency structure using 7 different frequencies from 500 Hz to 800 Hz in steps of 50 Hz. Within each stimulus train, all tones were of one frequency and were followed by a train of tones of a different frequency. The number of times the same tone was presented within one stimulus train varied pseudo-randomly between 1 and 11 (t = 1,…,11) such that 5% of all stimulus trains consisted of 1–2 identical tones, 7.5% of all stimulus trains consisted of 3–4 identical stimuli, and 87.5% of all stimulus trains consisted of 5–11 identical stimuli. For each participant and each session a different sequence of tones was generated online.

Following the suggestion that MMN assessment is optimal when the subject's attention is directed away from the auditory domain (4), subjects performed a distracting visual task and were instructed to ignore the sounds. Whenever a fixation cross changed its luminance, which occurred pseudo-randomly every 2–5 s (not coinciding with auditory changes), subjects had to press a button. One experimental session lasted approximately 15 min.

#### 1.3 Preprocessing of EEG data

Pre-processing and data analysis was performed using SPM12 (http://www.fil.ion.ucl.ac.uk/spm/). Continuous EEG recordings were referenced to the average, high-pass filtered using a Butterworth filter with cutoff frequency 0.5 Hz, down-sampled to 256 Hz, and low-pass filtered using Butterworth filter with cutoff frequency 30 Hz. The data were epoched into 500 ms segments around tone onsets, using a pre-stimulus baseline of 100 ms.

We rejected all trials overlapping with eye blink events, as detected by a thresholding routine on the vertical EOG channel, which was created from subtracting the activity of two additional electrodes which were attached infraorbitally and supraorbitally to the left eye. Finally, an artifact rejection procedure was applied using a thresholding approach on all EEG channels to detect problematic trials or channels. Trials in which the signal recorded at any of the channels exceeded 80 μV relative to the pre-stimulus baseline were removed from subsequent analysis, and channels in which more than 20 % of trials had to be rejected were marked as bad and subsequently interpolated for sensor-level statistics.

Bad channels occurred for three subjects in the placebo session (with 5, 1, and 1 bad channels, respectively) and for two subjects in the ketamine session (with 1 and 2 bad channels, respectively). Additionally, we had to mark two channels (F1 and C2) in five data sets as bad

(2 in placebo, 3 in ketamine) due to incorrect cabling. To exclude the possibility that our main results were driven by the interpolation of missing channel data, we performed all statistical analyses on the group level once without subject 14, who lost five channels due to bad signal quality, and once without the four subjects affected by the cabling errors. The results of the paired T-tests between the placebo and the ketamine session for these settings are reported in the supplementary tables S3 and S4.

#### 1.4 Computational Model

In the following, we provide more details on the perceptual model we used to simulate trial-by-trial estimated of prediction errors. We denote scalars by lower case italics (e.g., $x$), vectors by lower case bold letters (e.g., $\mathbf{x}$), and matrices by upper case bold letters (e.g., $\mathbf{X}$). Elements of vectors or matrices are referred to by subscripts (e.g. $X_{ij}$ denotes the $j_{th}$ element of the $i_{th}$ row of matrix $\mathbf{X}$). Trial numbers are indexed by the superscript $(k)$, e.g., $x^{(k)}$.

##### Perceptual Model: The Hierarchical Gaussian Filter (HGF)

In the HGF, sensory inputs are generated by a hierarchy of hidden states whose evolution in time is described as Gaussian random walks, where the step-size on each level is a function of the level above (Figure 1 of main paper). The uppermost level is assumed to evolve with a constant step-size, and the lowest level gives rise to the experimental stimuli the agent encounters. The variational inversion of this model under a mean field approximation yields simple one-step update equations that, collectively, represent a recognition model in which beliefs are updated by precision-weighted prediction errors (PEs).

Applying this to the present task, the agent infers on two hidden states of the world: a matrix $\mathbf{X}_2$ of transition probabilities (in logit space) between tones of different frequencies and the volatility $x_3$ (i.e., the degree to which the elements of $\mathbf{X}_2$ change from trial to trial).

On each trial, the elements of $\mathbf{X}_2$ determine the probability of encountering each of the possible transitions ($\mathbf{X}_1$), which in turn give rise to a particular tone being heard ($\mathbf{u}$). Because the matrix $\mathbf{X}_2$ encodes the logit-probability of transitions, where the column index indicates the previous tone and the row index the current one, only one column applies to each transition.

Given the straightforward discriminability of stimuli in our experiment, one can assume a simple binomial mapping from $\mathbf{X}_1$ to $\mathbf{u}$. Put differently, one can assume that any perceptual uncertainty about the tone category only arises from the probabilistic nature of the underlying cause, not from sensory noise:

$$p\left(x_{1,i,j}^{(k)}\right) = s\left(x_{2,i,j}^{(k)}\right)^{x_{1,i,j}^{(k)}} \left(1 - s\left(x_{2,i,j}^{(k)}\right)\right)^{1-x_{1,i,j}^{(k)}} \tag{1}$$

and

$$p\left(u_i^{(k)} \middle| x_{1,i,u^{(k-1)}}^{(k)}\right) = \left(u_i^{(k)}\right)^{x_{1,i,u^{(k-1)}}^{(k)}} \left(1 - u_i^{(k)}\right)^{1-x_{1,i,u^{(k-1)}}^{(k)}}, \tag{2}$$

where $s(x)$ is a logistic sigmoid (softmax) function:

$$s(x) \equiv \frac{1}{1 + \exp(-x)} . \tag{3}$$

The evolution of the elements of the transition matrix $\boldsymbol{X}_2$ in time are given by Gaussian random walks, where the step-size is a function of the current value of volatility $x_3$:

$$p\left(x_{2,i,j}^{(k)}\right) = \mathcal{N}(x_{2,i,j}^{(k)};\ x_{2,i,j}^{(k-1)}, \exp\left(\kappa x_3^{(k)} +\ \omega\right)) \tag{4}$$

Finally, $x_3$ evolves in time with a constant step-size $\vartheta$:

$$p\left(x_3^{(k)}\right) =\ \mathcal{N}\left(x_3^{(k)};\ x_3^{(k-1)}, \vartheta\right). \tag{5}$$

The three parameters governing this model, $\kappa$, $\omega$, and $\vartheta$, determine the strength of the coupling between the second and third level, the log-volatility (trial-by-trial variability) on the second level that is independent of the third level, and the variability of the volatility over time (meta-volatility), respectively. Here, we fixed the coupling parameter $\kappa$ to 1 because the scale of $x_3$ is arbitrary in our setting (for details, see (5)). This effectively eliminates this parameter from the model.

#### Inversion of the Model: The update equations

Inverting the above generative model yields posterior densities of (updated beliefs about) the hidden environmental states which caused the sensory inputs. These updated beliefs will be denoted in the following by their sufficient statistics (assuming Gaussian distributions), i.e., $\mu$ (mean) and $\sigma$ (variance/uncertainty) or the inverse of uncertainty, $\pi$ (precision/certainty; Figure 1 of main paper). Predictions, i.e., prior beliefs about the hidden states (before hearing the tone), will be denoted with a hat symbol (e.g., $\hat{\mu}$).

Using variational Bayes under a mean-field approximation and an approximation to the posterior energy function (6) leads to analytical trial-by-trial update equations, where belief updating rests on precision-weighted PEs, as described by equation (6) of the main paper. In the multivariate binary HGF with three levels, the specific update equations for $\boldsymbol{\mu}_2$ and $\mu_3$ are as follows:

$$\mu_{2,i,j}^{(k)} = \hat{\mu}_{2,i,j}^{(k)} + \varepsilon_{2,i,j}^{(k)} = \mu_{2,i,j}^{(k-1)} + \sigma_{2,i,j}^{(k)}\delta_{1,i,j}^{(k)} \tag{7}$$

and

$$\mu_3^{(k)} = \hat{\mu}_3^{(k)} + w_2^{(k)}\varepsilon_3^{(k)} = \mu_3^{(k-1)} + w_2^{(k)}\sum_{i=1}^{7}\frac{\hat{\pi}_{2,i,j}^{(k)}}{\pi_3^{(k)}}\delta_{2,i,j}^{(k)} \tag{8}$$

with the (unweighted) PEs

$$\delta_{1,i,j}^{(k)} = \mu_{1,i,j}^{(k)} - s\left(\mu_{2,i,j}^{(k-1)}\right) \tag{9}$$

and

$$\delta_{2,i,j}^{(k)} = \frac{\sigma_{2,i,j}^{(k)} + \left(\mu_{2,i,j}^{(k)} - \mu_{2,i,j}^{(k-1)}\right)^2}{\sigma_{2,i,j}^{(k-1)} + \exp\left(\kappa\mu_3^{(k-1)} + \omega\right)}, \tag{10}$$

and the additional weighting factor

$$w_2^{(k)} = \frac{\kappa}{2}\exp\left(\kappa\mu_3^{(k-1)} + \omega\right). \tag{11}$$

In these equations, the index *j* indicates the previously heard tone. As this is fixed for each trial, there are in total 7 updates on the second level per trial (*i* = 1,…,7), one for each possible tone. For $\varepsilon_2$, we chose the index *i* equal to the index of the actual presented tone of the current trial (see below).

For a detailed derivation of these equations and the updates of the precisions, and an interpretation in terms of precision-weighted PEs, the interested reader is referred to (6). These updates provide approximately Bayes-optimal rules for the trial-by-trial updating of the agent's beliefs about the probability of encountering a tone of a given frequency on the next trial. Note that this Bayes-optimality is individualized in the sense that it is optimal with respect to given values of the model parameters and the starting values of beliefs and belief precisions, which may differ across subjects.

In our context, because the auditory MMN is a passive paradigm (no behavior in response to the tones is recorded; subjects are instructed to ignore the sounds), we cannot infer the values of these parameters from behavioral responses (5). Therefore, given the subject- and session-specific stimulus sequences, we simulated belief trajectories that are expected under Bayes-optimal parameter values (defined as the parameter values that result in minimal overall surprise about the stimulus sequence encountered). This results in slightly different parameter values for each subject and each session because the tone sequences differed slightly (they were generated, under identical probabilities across subjects, on the spot during each session). The parameter estimates and the priors used are summarized in supplementary Tables S1 and S2. There were no significant differences in these parameter values between placebo and ketamine conditions.

| | Prior mean | Prior variance |
|---|---:|---:|
| $\kappa$ | $1$ | $0$ |
| $\omega$ | $-6$ | $25$ |
| $\vartheta$ | $0.05$ | $0.088$ |
| $\mu_3^{(0)}$ | $1$ | $0$ |
| $\sigma_3^{(0)}$ | $\log(0.1)$ | $1$ |
| $\mu_{2ij}^{(0)}$ | $\mathrm{logit}(\frac{1}{49})$ | $0$ |
| $\sigma_{2ij}^{(0)}$ | $\log(1)$ | $0$ |

**Table S1:** Priors on HGF perceptual parameters and starting values. All parameters and starting values were fixed (i.e., not estimated) except for the tonic learning rates $\omega$ and $\vartheta$, and the starting value of the high-level uncertainty $\sigma_3^{(0)}$.

| | Placebo | | Ketamine | | Diff.(t) |
|---|---|---|---|---|---|
| | Mean | Std. | Mean | Std. | $p$ |
| $\omega$ | $-10.04$ | 0.20 | $-10.06$ | 0.28 | 0.83 |
| $\vartheta$ | 0.045 | 0.003 | 0.046 | 0.003 | 0.33 |
| $\sigma_3^{(0)}$ | 0.1001 | 2.1e−04 | 0.1001 | 3.8e−04 | 0.67 |

**Table S2:** Average posterior means of estimated parameter values for surprise-minimizing agents across subjects and sessions. There were no significant differences between the parameters in the placebo and the ketamine sessions.

#### The computational quantities of interest: precision-weighted prediction errors

The precision-weighted PE about transition identity ($\varepsilon_{2,i,j}^{(k)}$) is a low-level PE in that it refers to the stimulus outcome (first level of the model) and serves to update the estimate of the transition matrix (second level). Out of the seven updates on each trial (one for each tone category), we included the transition PE, i.e., the PE on the transition that actually occurred on a given trial, which is by definition always a positive PE.

The precision-weighted PE about the transition matrix $\varepsilon_3^{(k)}$ is a high-level PE that refers to transition probabilities (second level of the model) and serves to update the (log) volatility estimate $\mu_3^{(k)}$ at the highest level of the model. This PE is, on each trial, proportional to the sum over all 7 updates in the transition matrix, because all entries in the matrix share a common volatility estimate on the third level. It is a signed PE, because the (log) volatility is a continuous quantity.

Supplementary Figure S1 shows the simulation results for an example subject and session, including the trajectories of precision-weighted PEs on both levels. The corresponding parametric modulators in the GLM were modelled as events that were time-locked to the tone onset in each trial. At first glance, it may seem that the $\varepsilon_3$regressor simply amounts to a drift-like signal (Figure S1B). This, however, is not the case; the design of our experiment, with prolonged trains of identical stimuli that exchange each other, leads to separate monotonic changes in log-volatility estimates for standard and deviant trials, with jump-like transitions between them (Figure S1C).

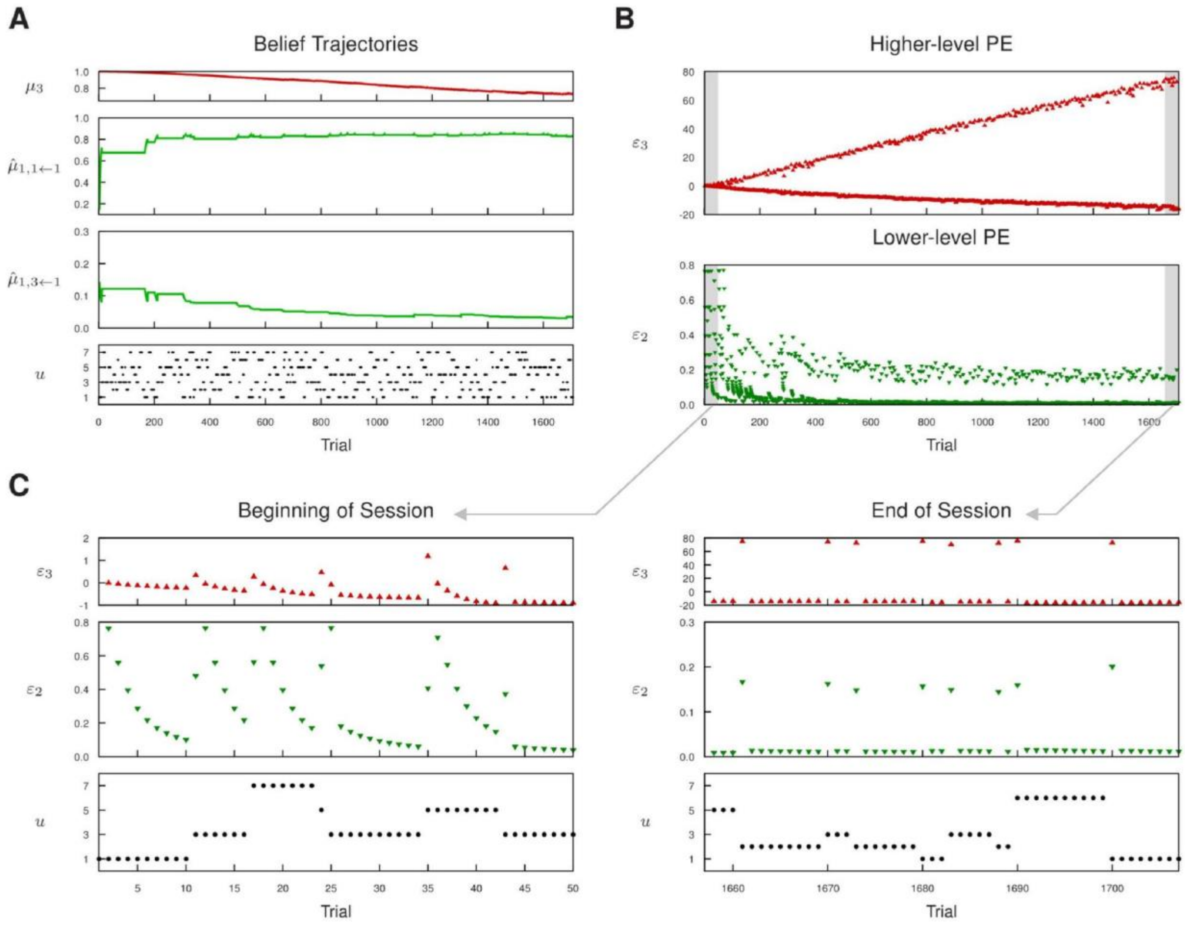


**Figure S1.** Simulation Results: Example Trajectories from a representative subject in the placebo session. **A** Time course over the experimental session (1707 trials) for the agent's simulated belief about volatility ($\mu_3$, in red), its trial-by-trial prediction of the repetition probability of tone 1 ($\hat{\mu}_{1,1,1}$, in green), its trial-by-trial prediction of the transition probability from tone 1 to tone 3 ($\hat{\mu}_{1,3,1}$, in green), and the tone sequence it was exposed to ($u$, black dots). After an initial learning period, the agent has learned that repetitions are likely, and transitions are unlikely. **B** Trial-by-trial values of the two precision-weighted PEs used as regressors in the GLM of the EEG signal. The initial learning period is characterized by high low-level PEs about stimulus occurrences ($\varepsilon_2$) for all tone events, later the deviant trials (tone transitions) separate clearly from the standard trials (repetitions). As the agent's volatility estimate decreases, the higher-level PE ($\varepsilon_3$) in response to deviants separates more and more from the one in standard trials. **C** The two PE trajectories in more detail for the 50 first tones of the session (left) and the 50 last tones of the session (right). At first, all tones are similarly surprising ($\varepsilon_2$ in the beginning of the session), but in the end, because the agent's estimates of repetition and transition probabilities are stable and accurate, repetitions elicit almost no PEs anymore.

#### 2 Supplementary Results

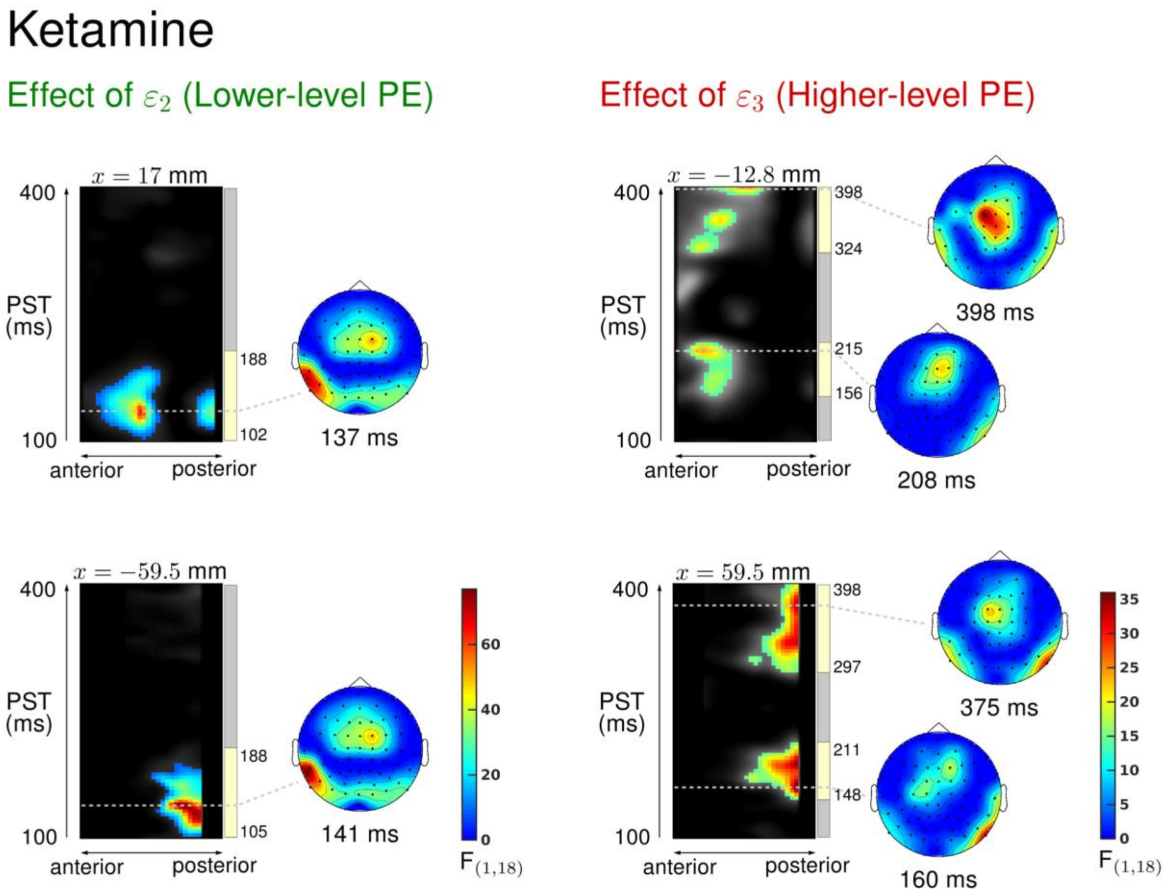


**Figure S2:** Results of the model-based EEG analysis in the ketamine condition: effects of the high- and the low-level PE. The left side always shows an F-map across the scalp dimension *y* (from posterior to anterior, x-axis), and across peristimulus time (y-axis), at the spatial *x*-location indicated above the map. Significant F values ($p < 0.05$, whole-volume FWE-corrected at the cluster-level with a cluster-defining threshold of $p < 0.001$) are color-coded according to the legend plotted on the right. Time-windows of significant correlations are indicated by the yellow bars next to the colored clusters of significant F values. The scalp maps next to the F-maps always show the F-map at the indicated peristimulus time point, corresponding to the peak of that cluster, across a 2D representation of the sensor layout. We found significant correlations of the EEG signal with our two computational quantities across fronto-central and temporal channels. For the lower-level PE, $\varepsilon_2$, the correlation peaked at 137 ms at fronto-central channels, and at 141 ms at left temporal channels. For the higher-level PE, $\varepsilon_3$, it peaked at 208 ms and at 398 ms after stimulus onset at fronto-central channels, and at 160 ms and 375 ms at right temporal channels.

| Higher-level PE ($\varepsilon_3$) | Peak Coordinates | | | peak-level | | | cluster-level | |
|---|---|---|---|---|---|---|---|---|
| | $x$ [mm] | $y$ [mm] | $t$ [ms] | $T_{\{17\}}$ | $Z_{\equiv}$ | $p_{\text{FWE}}$ | $k_{\text{E}}$ | $p_{\text{FWE}}$ |
| Placebo>Ketamine | $-4$ | 2 | 223 | 5.57 | 4.14 | 0.061 | 564 | 0.009 |

**Table S3:** Significant drug differences for the representation of the higher-level PE $\varepsilon_3$ in Placebo vs. Ketamine condition, when excluding subject 14 (for whom five channels had to be rejected due to noise, resulting N=18). The table lists the peak coordinates, t values, corresponding Z values, FWE-corrected p-values at the voxel level, cluster size ($k_E$) and whole-volume FWE-corrected p-values at the cluster level of voxels showing significantly stronger representation of $\varepsilon_3$ under placebo compared to the ketamine condition (paired t-test, $p < 0.05$ whole-volume FWE-corrected at the cluster-level with a cluster-defining threshold of $p < 0.001$). No significant drug differences were found for the lower-level PE.

| Higher-level PE ($\varepsilon_3$) | Peak Coordinates | | | peak-level | | | cluster-level | |
|---|---|---|---|---|---|---|---|---|
| | $x$ [mm] | $y$ [mm] | $t$ [ms] | $T_{\{14\}}$ | $Z_{\equiv}$ | $p_{\text{FWE}}$ | $k_{\text{E}}$ | $p_{\text{FWE}}$ |
| Placebo>Ketamine | $-4$ | 2 | 227 | 5.69 | 4.03 | 0.099 | 442 | 0.035 |
| | 8 | 13 | 230 | 5.56 | 3.97 | 0.117 | | |
| | $-34$ | $-14$ | 230 | 4.18 | 3.31 | 0.528 | | |

**Table S4:** Significant drug differences for the representation of the higher-level PE $\varepsilon_3$ in Placebo vs. Ketamine condition, when excluding the four subjects affected by cabling errors (see main text, resulting N=15). The table lists the peak coordinates, t values, corresponding Z values, FWE-corrected p-values at the voxel level, cluster size ($k_E$) and whole-volume FWE-corrected p-values at the cluster level of voxels showing significantly stronger representation of $\varepsilon_3$ under placebo compared to the ketamine condition (paired t-test, $p < 0.05$ whole-volume FWE-corrected at the cluster-level with a cluster-defining threshold of $p < 0.001$). No significant drug differences were found for the lower-level PE.
